# Precursor types predict the stability of neuronal branches

**DOI:** 10.1101/2021.04.23.441127

**Authors:** Joachim Fuchs, Britta J. Eickholt

## Abstract

Branches are critical for neuron function, generating the morphological complexity required for functional networks. They emerge from different, well-described, cytoskeletal precursor structures that elongate to branches. While branches are thought to be maintained by shared cytoskeletal regulators, our data from mouse hippocampal neurons indicate that the precursor structures trigger alternative branch maintenance mechanisms with differing stabilities. While branches originating from lamellipodia or growth cone splitting events collapse soon after formation, branches emerging from filopodia persist. Furthermore, compared to other developing neurites, axons stabilise all branches and preferentially initiate branches from filopodia. These differences explain the decreased stability of branches we observe in neurons lacking the plasma membrane protein phospholipid phosphatase related protein 3 (PLPPR3/PRG2). Rather than altering branch stability directly, PLPPR3 boosts a ‘filopodia branch program’ on axons, thereby indirectly initiating more long-lived branches. We propose that studies on branching should distinguish overall stabilising effects from effects on precursor types, ideally using multifactorial statistical models as exemplified in this study.

## Introduction

Rivers, lightnings or trees - branches are ubiquitous in both receptive and transmitting processes in nature. Their abundance ensures optimal coverage of area, balancing maximal receptivity with the shortest distance to their origin. It is no surprise that also neurons use branches to optimise their signalling efficacy in a neuronal network. The cellular mechanisms controlling this specific branching behaviour have been studied for 30 years (summarised in Kalil and Dent, 2014). Recently, branching gained attention in the setting of regenerative growth of central nerve cells (Griffin and Bradke, 2020). Strategies to improve recovery following injury to the CNS include promoting branch formation in non injured neurons to form alternative pathways (Fink et al., 2017) or inhibiting branching in injured neurons to facilitate undisturbed elongation (Tedeschi et al., 2019).

The main drivers of the remarkable morphology of neurons, the actin and microtubule components of the cytoskeleton, participate sequentially in branch formation. Initially, F-actin structures – filopodia or lamellipodia – remodel the plasma membrane. Subsequently, de-bundling, transport, and polymerisation of microtubule arrays into the actin-enriched protrusion elongate the emerging branch. Not every branch will persist, however. Branches are maintained mainly by stabilising the microtubule cytoskeleton (Gallo, 2011; Kalil and Dent, 2014). Supporting this sequential model, F-actin regulators have been described to control branch emergence, while branch elongation requires crosslinkers between the two cytoskeletal structures, and branch maintenance is influenced predominantly by microtubule-associated proteins (Armijo-Weingart and Gallo, 2016).

Neurons establish networks by connecting selectively to specific regions and by stablishing layer specific receptive fields (Cuntz et al., 2010). Especially in long axons, this requires preventing most branching (Gibson and Ma, 2011). To circumvent the suppression of branching at specific sites, neurons rely on several extrinsic and intrinsic cues. One major regulator of branching, the PI3K/PTEN pathway, triggers F-actin accumulation and protrusion formation (Kakumoto and Nakata, 2013; Ketschek and Gallo, 2010), induces transport and local translation (Akiyama and Kamiguchi, 2010; Spillane et al., 2013), and regulates microtubule stability (Kath et al., 2018). We previously described a transmembrane protein, plasticity related gene 2/phospholipid phosphatase related protein 3 (PRG2/PLPPR3), that is able to relieve the general branch suppression by inhibiting PTEN, the negative regulator of PI3K signalling. Specifically, PLPPR3 redistributes growth towards branches by inducing filopodia formation (Brosig et al., 2019). Given that PI3K signalling is involved in multiple steps of branching, we set out to analyse the contribution of PLPPR3 to later stages, specifically to branch maintenance.

### Plppr3^−/−^ branches are less stable

We employed phase-contrast microscopy of cultured mouse neurons from wildtype (*WT*) and *Plppr3*^−/−^ hippocampus. Neurons were imaged for 24 h, at a temporal precision of 10-min intervals. In initial observations, *Plppr3^−/−^* branches appeared less stable (Supplementary movie 1). We therefore measured the lifetime of each branch as the difference between the time of initiation and, if applicable, collapse, in a blinded and randomised manner. When comparing these raw lifetimes of *WT* and *Plppr3*^−/−^ branches, the difference between the two genotypes is too small to be statistically detectable (p=0.83, Figure 1A).

**Figure 1:**
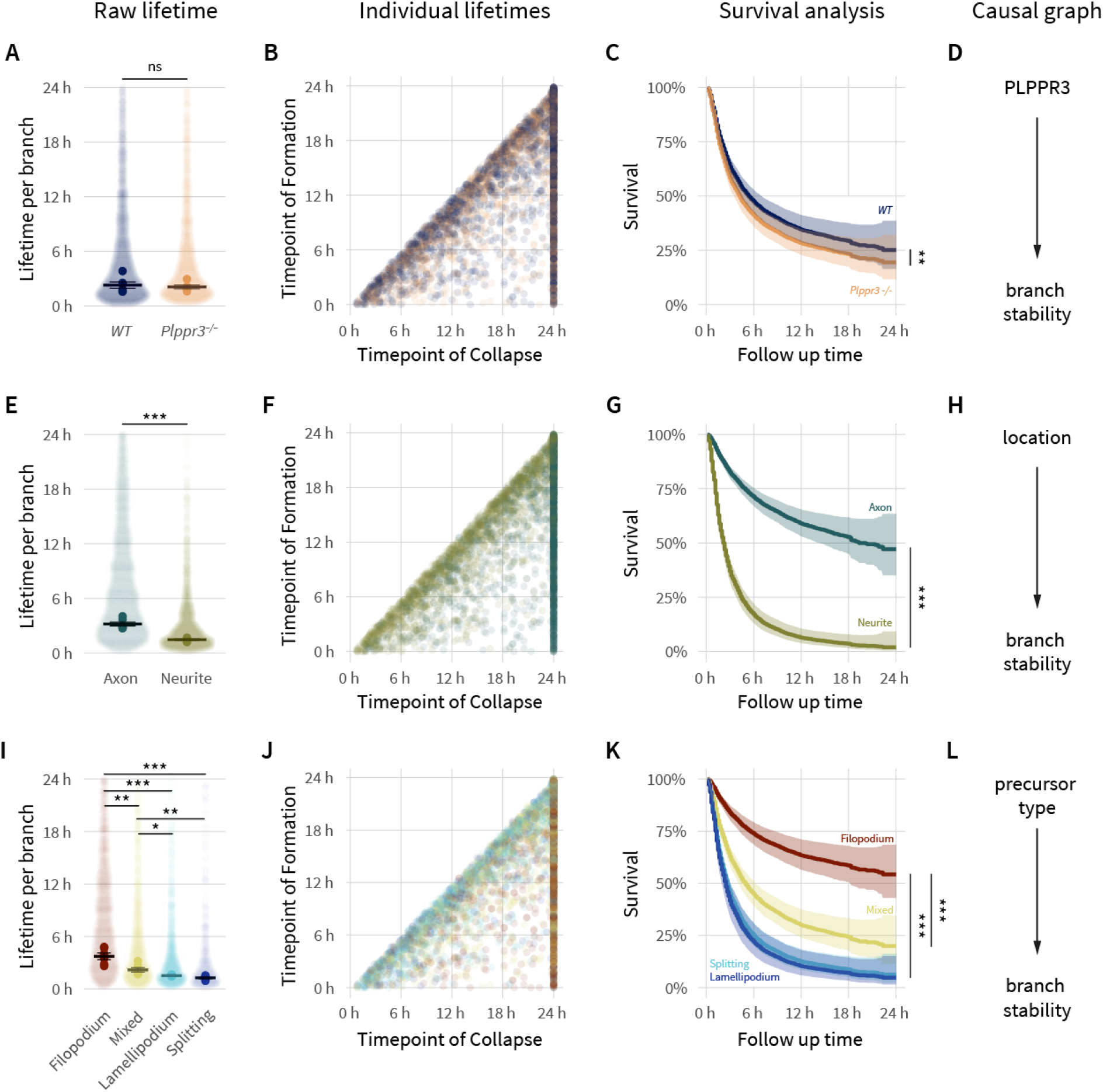
Individual effects on neuronal branch stability. *(A) Raw lifetimes of individual branches (opaque dots) and per experiment average (solid dots) show a small, statistically non-detectable effect of PLPPR3 loss. (B) Scatterplot of timepoint of formation versus collapse for individual branches shows strong right-censoring of the data. Note the similar overall distribution of WT and Plppr3^−/−^ branches. (C) Correcting for censoring bias using survival analysis reveals a small but statistically detectable effect of Plppr3-loss on branch lifetime. (D) Causal graph highlighting connection of PLPPR3 to branch stability. Panels E-H show corresponding graphs for axonal vs. non-axonal branches. (G) Survival analysis shows a strong effect of axonal location on branch stability in both WT and Plppr3^−/−^. (H) Causal graph highlighting connection of location to branch stability. Panels I-K show the corresponding graphs for the different branch precursor types. (K) Survival analysis shows the strong effect of the branch precursor type on stability in both WT and Plppr3^−/−^, with filopodia initiating the most stable branches. (L) Causal graph highlighting connection of precursor type to branch stability. n_Ind_=2317 (WT) & 2165 (Plppr3^−/−^), n_Exp_=6 (six independent cultures), transparent ribbons show 95% confidence intervals in survival curves, error bars in the raw lifetime plots indicate SEM between experiments. * p<0.05, ** p<0.01, *** p<0.001*

A drawback of such analyses is that the raw lifetime estimate is extremely biased towards short lifetimes. In our experiments, 40% of the branches did not collapse during the observed time window (heavily right-censored data, Figure 1B). We therefore analysed the data with a standard method in clinical trials developed to handle censored datasets, survival analysis, also termed time-to-event analysis. This established the *Plppr3* genotype as a predictor for branch stability (Figure 1C&D, hazard ratio (HR): 1.2 (95% CI: 1.1-1.3), p=0.006). Counterintuitively however, this effect is not evidence that PLPPR3 *directly* affects branch maintenance. Deeper analyses establish the effect of PLPPR3 on branch stability as secondary to its primary effect of inducing one specific branch precursor, filopodia.

### Branch precursor and location predict branch stability

During our analyses of branch stability in developing neurons, axonal branches appeared to persist longer than branches on immature dendrites, hereafter termed neurites (Supplementary movie 1). Supporting this, survival analyses of these branches distinguishing their location show a strong risk of collapse for branches on neurites (Figure 1E-G, HR=5.2 (4.6-5.9), p<0.001). The stabilising effect on developing axons is likely influenced by differences in orientation and post-translational modifications of microtubules (reviewed in Janke and Magiera, 2020) and the contribution of distinct microtubule binding proteins in axons (reviewed in Bodakuntla et al., 2019; and Conde and Cáceres, 2009). The observed increased stability for axon branches is likely controlled by the same mechanisms that govern microtubule stability in the axon (Figure 1 H).

To our surprise, however, the type of actin precursor *also* strongly influenced subsequent branch stability (Figure 1I-K). We classified precursor types by morphology (Supplementary figure 1) as bifurcations of the growth cone (‘splitting’) or as formations on the axon shaft (collateral branches) originating from thin filopodia or sheet-like lamellipodia. With many collateral branches initiating from filopodia being invaded by lamellipodia directly before branch elongation (Flynn et al., 2009; Withers and Wallace, 2020), we added a hybrid class (‘mixed’). Our quantification revealed that branches originating from lamellipodia and growth cone splitting were at a high risk of collapse within a few hours (HR_Spl_=5.0 (3.8-6.6), HR_Lam_=4.6 (3.6-5.7), both p<0.001), while most branches originating from filopodia remained stable throughout the 24h imaging period. Mixed precursor branches were at an intermediate risk of collapse (HR_Mix_=2.6 (2.1-3.3), p<0.001). While lamellipodial branches appear to be stochastic “trial-and-error” branch initiations, filopodial branches may be a deterministic mode that form to stay.

### Axons initiate similar numbers of branches but use different precursors than neurites

Similar to previous studies (Armijo-Weingart and Gallo, 2016), neurons in our experimental setup initiate most branches as collaterals, with comparable numbers of lamellipodial, filopodial and ‘mixed’ precursor branches, while bifurcations are scarce (Figure 2A&C). Furthermore, while the total number of branch initiation events is similar between axons and neurites (Figure 2A), the proportions of the precursor types differ drastically, with filopodia mainly dominating axon branch initiations and lamellipodia being the predominant neurite branch precursor. This difference becomes even more apparent over time, when visualising which precursor type the branches originate from (Figure 2B). Most branches on axons originate from filopodia and mixed precursors and accumulate fast, while branches on early neurites hardly accumulate, irrespective of the precursor type. In addition to using the most efficient precursors, axons therefore also seem to stabilise branches of all precursors.

**Figure 2:**
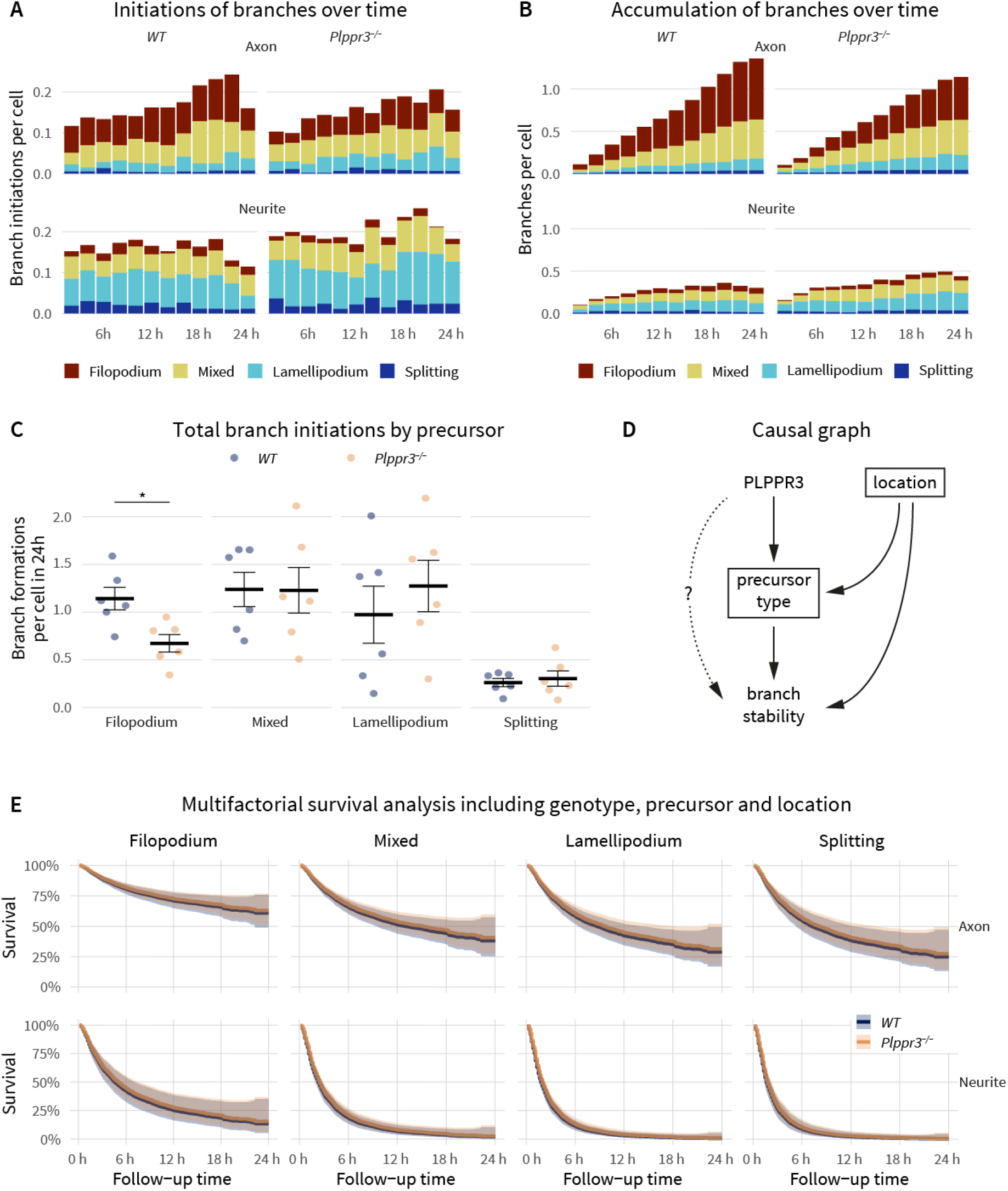
Combined influences on branch stability. *Distribution of branch forming events over time, colour-coded by precursor type, according to location as well as genotype. Note the similar overall number of branch initiations but different composition of precursor types between axons and neurites. Branches per cell present over time, colour-coded by precursor type, according to location as well as genotype. Note the increased branch accumulation on axons vs neurites and the different precursor composition between branch initiations (panel A) and accumulating branches (panel B). (C) Total branch formations per cell over 24 h, originating from different precursor types and genotype. Plppr3^−/−^ neurons initiate fewer branches from filopodia. (D) Directed acyclic graph aggregating the information from Figures 1 & 2. According to causal graph theory, to quantify a direct effect of PLPPR3 on branch stability in an unbiased way, a multifactorial analysis has to account for precursor type as well as location (indicated by squares). (E) Survival analysis of WT vs. Plppr3^−/−^ branches including location as well as precursor types as covariates demonstrates no direct effect of PLPPR3 on branch stability. Forest plot of model: Supplementary figure 3. n_Ind_=2317 (WT) & 2165 (Plppr3^−/−^), n_Exp_=6. The opaque ribbon in (E) shows the 95% confidence interval of the survival curves. Error bars in (C) indicate SEM between experiments. * p<0.05*

Compared to *WT*, *Plppr3^−/−^* neurons initiate fewer branches from filopodia precursors (Figure 2A&C, p=0.046), without affecting other precursor types. *Plppr3^−/−^* neurons also seem to accumulate fewer branches on axons (Figure 2B). This is in accordance with our recent study indicating a lower number of filopodia in *Plppr3*^−/−^ specifically on early polarised axons, and a lower number of axon branches at later developmental stages. Notably, the effect sizes of filopodia density, branch density (Figure 6 in Brosig et al., 2019) and branch initiations from filopodia (this study) are very similar. This indicates that the initial filopodia number is the main determinant of the branching defect in *Plppr3^−/−^* neurons rather than a defect in filopodia-to-branch transitions.

### PLPPR3 regulates branch stability via the precursor type alone

We showed that PLPPR3-loss decreases branch stability and reduces the numbers of the most efficient precursor (filopodia), preferentially on axons, which themselves stabilise branches in two distinct ways. With such individual but interdependent information it is difficult to distinguish whether the PLPPR3 effect on branch stability is direct, as is conceivable by inducing PI3K signalling and microtubule stability, or indirect, by regulating the number of the more efficient filopodia precursors.

This information, however, helps to generate an informed causal diagram (directed acyclic graph [DAG], (Pearl, 1995), which in turn can form the basis of a multifactorial statistical analysis quantifying the contribution of the various effects. The causal diagram for these data (Figure 2D) assumes that both precursor type and location directly influence branch stability, and PLPPR3 directly influences the abundance of one precursor type. Furthermore, the location alters the distribution of precursor types. According to causal graph theory (Pearl et al., 2016), to determine whether there is a direct effect of PLPPR3 on branch stability (dotted line in Figure 2D), the influence of the precursor type as well as location (precursor type being a collider) have to be accounted for to obtain an unbiased estimate.

When adjusting a survival analysis for the contribution of precursor type and location, the stability of *Plppr3^−/−^* branches is indistinguishable from WT branches (Figure 2E & Supplementary figure 3, HR=0.9 (0.8-1.1), p=0.26). This still means that *Plppr3^−/−^* branches are less stable than WT (Figure 1C), however there is no evidence in this dataset for a *direct* effect of PLPPR3 on branch stability. The effect of PLPPR3 on branch stability is fully explainable as an *indirect* consequence of its effect on filopodia abundance.

### Branch precursor types initiate distinct branching systems

Our analysis of branch stability in hippocampal neurons highlights a strong influence of location as well as precursor type. We identified that filopodia are not the most abundant precursor, but the most efficient. Losing filopodia branches (in *Plppr3*^−/−^ neurons) appears to decrease the stability of the remaining branches, while a recent study suggested that specifically losing lamellipodia trended towards increased branch stability in a sample of only three neurons (Pollitt et al., 2020). Both treatments appear to shift the equilibrium of branch precursor types, without affecting the stability of branches from each individual precursor. This suggests the different precursors initiate distinct types of branches with mechanistically independent maintenance programs.

This raises the question of how the configuration of the actin cytoskeleton in the precursor type influences branch stability hours after the precursor structures elongate? The mechanical rigidity of parallel-bundled F-actin in filopodia and the meshwork of F-actin in lamellipodia may account for differences in branch stability, as forces generated by filopodia and lamellipodia in growth cones differ (Cojoc et al., 2007), and microtubule growth is receptive to force (Janson et al., 2003). Alternatively, branch precursors may recruit different microtubule crosslinkers (Dogterom and Koenderink, 2019) for microtubule capture in forming branches, or use different mechanisms to supply fresh microtubules via severing, transport or de novo nucleation. Differential modes of branch initiation have been described for the microtubule severing-enzymes katanin and spastin (Yu et al., 2008). Most importantly, branches from different precursors can be expected to differ in the microtubule stabilising factors they recruit.

Microtubules and their binding proteins, however, can also actively modify the actin precursors. Lamellipodial actin waves both co-occur with and require dynamic microtubules (Winans et al., 2016). The microtubule-binding proteins doublecortin (Fu et al., 2013) and Gas2L1 (Willige et al., 2019) regulate F-actin stability, altering axon branching. Reduction of Map7 induces more branch initiations while decreasing branch stability (Tymanskyj et al., 2017), which might be explained by inducing more lamellipodia branches or by separate effects on microtubule stability and precursor types. Given that the developing axon seems to preferentially use efficient precursors *and* to stabilise branches in general, future studies on branch stabilising factors should distinguish effects on all branches from those on specific precursors.

### Cell biology can benefit from multifactorial analyses informed by causal models

In addition to these biological findings, this study highlights how interpreting the effects of multiple interdependent factors *independently* can misinform the mechanistic models inferred from data, often establishing more pathways than the data accounts for. While cell biology routinely uses statistical tests to protect against false positive findings in individual experiments, the integration of evidence from multiple sources by collecting and discussing data (e.g., in scientific reviews) does not formally test their relationship. Consequentially, resulting models often contain more connections than are experimentally verified.

Our work highlights the value of quantifying the relationship of individually published links in additional, multifactorial experiments. Fortunately, statistical (multiple regression) and causal tools (DAGs, counterfactuals) have evolved considerably recently (Hernán and Robins, 2020; Rohrer, 2018; Suttorp et al., 2015). They are frequently employed in other fields such as epidemiology (Greenland et al., 1999) or ecology (Greenacre and Primicerio, 2013; James and McCulloch, 1990) to reduce the number of false positive links in multifactorial systems. Our study exemplifies how multifactorial statistical analyses informed by causal graphs can advance cell biology, offering a clear benefit over unifactorial ANOVAs and t-tests, and leveraging this methodological approach has the potential to clarify the structure of biological pathways.

## Acknowledgements

We thank Kerstin Schlawe, Kristin Lehmann and Beate Diemer for excellent technical assistance. Funding was provided by the DFG SFB 958, A16 and SFB TRR 186, A10.

## Author contributions

B.J.E. and J.F. designed the experiments, J.F. performed experiments and analysis, and B.J.E. provided supervision and funding. Both authors wrote the manuscript and revised the final version.

## Methods

The experimental strategy is summarised in Supplementary figure 2, and a detailed description of each step is provided below.

## Animal procedures and primary neuron culture

Mice were housed and handled according to local ethical guidelines and approved animal care protocols (under the license T0347/11, Landesamt für Gesundheit und Soziales Berlin) according to the guidelines of the animal welfare of Charité Universitätsmedizin Berlin. The mice were housed in standardised conditions under 12-hour day-night cycle with water and food available ad libitum.

The *Plppr3^−/−^* line (described in Brosig et al., 2019) is maintained in a C57Bl/6 NCrl background. Heterozygous parents were bred for primary neuron culture preparation from day 16.5 embryos. Briefly, hippocampi of homozygous littermates (wildtype or knockout) were pooled for further single cell isolation without stratifying by sex. Extracellular matrix was degraded for 15 min with 10% trypsin in HBSS (Life Technologies), washed with HBSS with 1% horse serum and Neurobasal A medium before trituration to single cells with glass pipets.

Four-well glass-bottom chamber slides (μ-Slide, Ibidi) were coated sequentially with laminin (20 mg/ml) and poly-ornithine (15 mg/ml) before plating hippocampal neurons at a density of 25.000 / cm^2^ in Neurobasal A medium containing 2% B27 (Life Technologies), 1% penicillin/streptomycin (Life Technologies), 100 mM β-mercaptoethanol (Applichem) and 1% GlutaMAX (Life Technologies). Neurons were grown at 37 °C and 5% CO_2_ in a humidified incubator for 48 h before starting live cell imaging. Each individual neuron culture was considered as an independent N. The sample size was not estimated via power analysis prior to experiments due to lack of effect size estimates for the question under study. Instead, we chose the sample size to exceed typical sample sizes in cell biological experiments.

## Long-term live cell microscopy

Long-term live cell recordings were undertaken with a Nikon Eclipse Ti microscope equipped with a small stagetop as well as full incubator enclosure, to maintain cells at 37 °C (in the full incubator) and 5% CO_2_ and humidity (in the stagetop) throughout the imaging session. Growing hippocampal neurons were visualised using Köhler adjusted phase contrast (Ph2) brightfield microscopy in areas of similar density across conditions using Nikon’s Perfect Focus System to adjust for thermal fluctuations in focus. Three fields of view per genotype per litter were imaged for 24 h in 10-min intervals resulting in 18 movies from 6 cultures per group.

## Manual classification of branch events

Prior to analysis, movies were randomised and renamed automatically using a custom R script (Randomize_folder.R, https://github.com/jo-fuchs/Microscopy_analysis_snippets/tree/master/R) to perform the analysis blind to genotypes. For each movie, the total number of cells was recorded to adjust for differences in density between cultures. At each timepoint, newly forming branches (defined as processes longer than 10 μm) were marked with a region-of-interest (ROI) overlay using ImageJ/FIJI’s ROI-manager (Schindelin et al., 2012). The line colour of the ROI was encoded to represent four morphologically distinct precursor types – growth cone splitting (blue), filopodium (red), lamellipodium/actin wave (green) and a mixed type (yellow), see Supplementary figure 1.

In a second round of analysis, branch type classifications were quality controlled and for each branching event the location of formation was recorded as: (1) on a *neurite* of both polarised and non polarised neurons or (2) on the *axon* of clearly polarised neurons, defined as the persistently longest process of a neuron. Branching events on processes with cell bodies outside of the field of view were classified as *unclear* location. Additionally, the timepoint of collapse for each branch was recorded as the time at which a branch – or the originating process – completely retracted. The timepoint of collapse for branches that did not collapse during the recordings was set to the last frame and treated as censored data in subsequent analyses. All ROIs were saved and exported as comma separated value (csv) files named correspondingly to the movie.

## Statistical analysis of branch initiation and lifetime

All further analysis steps were performed in R/RStudio (R Core Team, 2020) and are fully documented at https://github.com/jo-fuchs/Branch-Lifetime-PRG2. Briefly, individual branch event and cell count files were merged and unblinded (merge_data.R) before merging the cell count with the individual branching data. Further cleaning steps (clean_data.R) included converting frame counts to hours, calculating lifetime, creating a censoring indicator (whether a branch was present in the last frame) and calculating inverse probability weights (1 / time until end of movie) to correct for the higher risk of censoring for branches forming close to the end of the data acquisition.

All statistical analyses are described in the R-scripts creating the figures (Figure_1.R and Figure_2.R). Lifetimes (Figure 1A, E & I) and branch formations per cell (Figure 2C) were summarised per experiment, assumptions for linear models were tested graphically using Residual vs Fitted, Q-Q, Scale Location and Cook’s distance plots. Due to heterogeneous variances, Welch’s t-test was used for pairwise comparisons. In cases of more than two comparisons, p-values were adjusted using Holm’s correction.

For the branch incidence (Figure 2A) and branch accumulation (Figure 2B) over time, branching and collapse events were binned into 2-h slots (cumulative_branches.R). For each bin, the net “flux” of branches (formed branches minus collapsed branches) was determined stratified by location, precursor type and genotype. The accumulation of branches was determined by the cumulative sum of this “flux”, normalised by the number of cells the branches originated from.

Survival analyses of branch lifetimes were computed using Cox proportional hazards models including individual factors (Figure 1C, G, K) or the full model presented in Figure 2D. Assumptions were tested using Schoenfeld’s test and inspecting residual plots, all models were weighted by the inverse probability weight calculated above, to account for the higher chance of censoring for branches forming close to the end of the imaging session. The final model was visualised in a forest plot (Supplementary figure 3). Final styling of the figures was performed in Adobe Illustrator 2021.

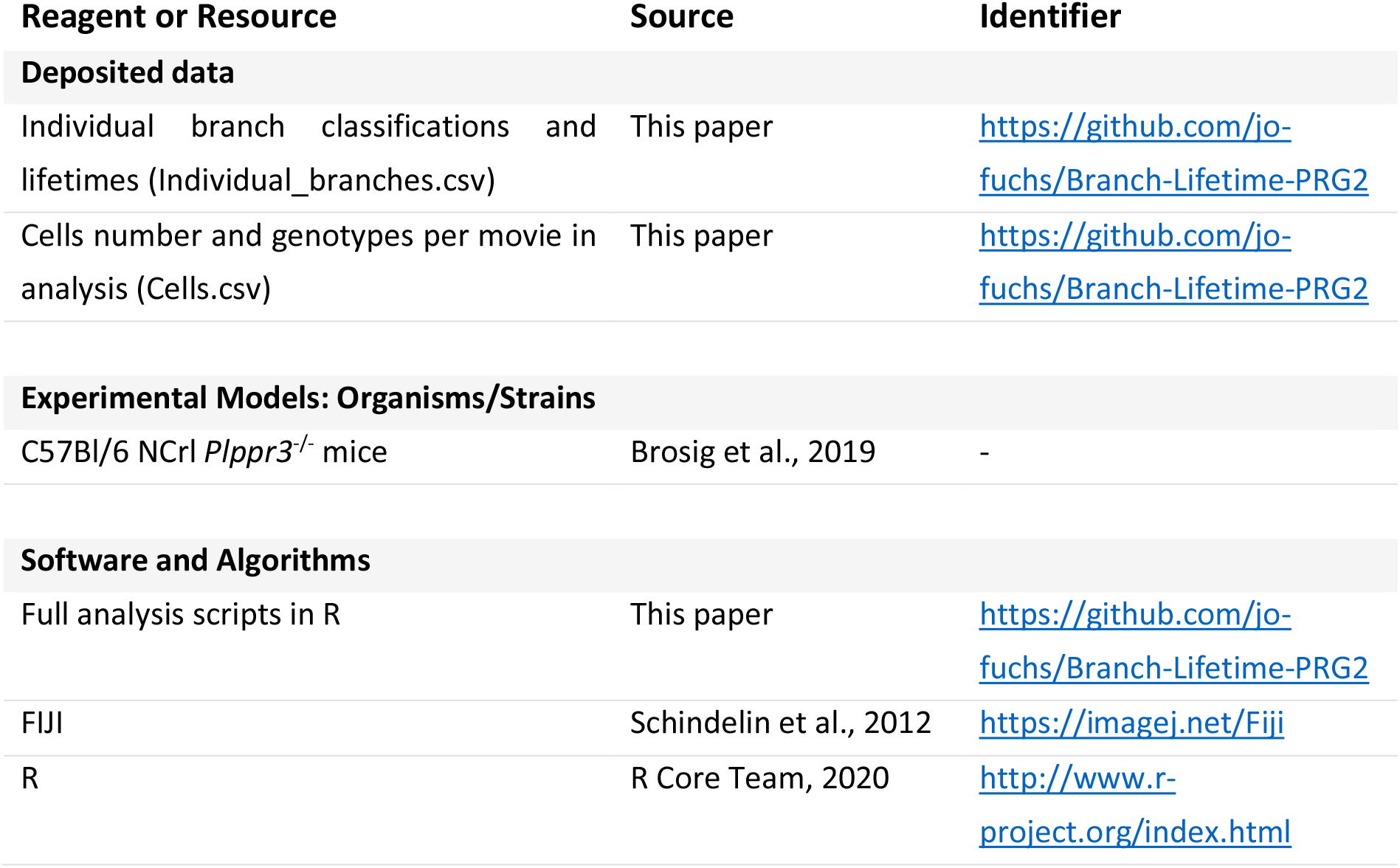

## Print Supplement: Example movies of WT and Plppr3^−/−^ branching behaviour

**Supplementary movie 1:**
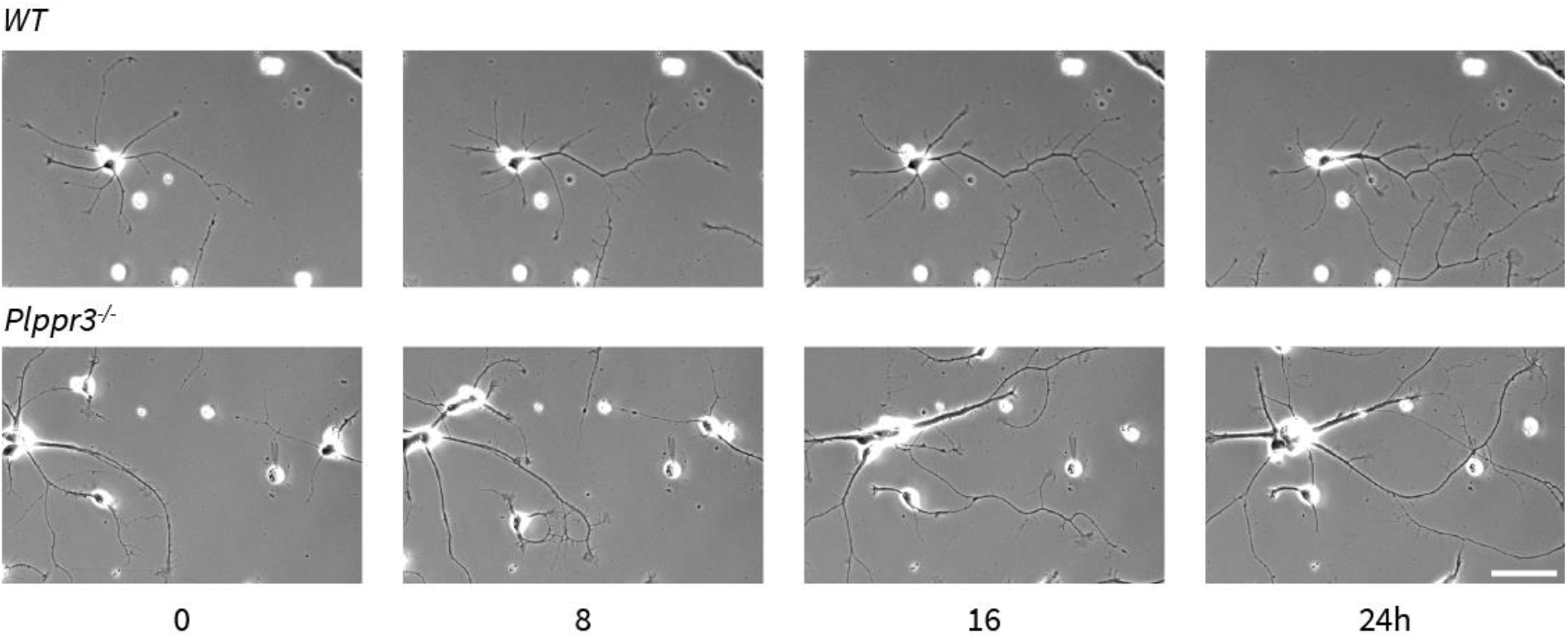
Timeseries of example movies of WT and Plppr3^−/−^ mouse hippocampal neurons. *WT neurons initiate many branches on both axons and neurites, but it was mainly axon branches that persisted. The majority of branches on neurites originated from lamellipodia, whereas the majority on axons were from filopodia. In Plppr3^−/−^ cells, fewer branches originated from filopodia, many short-lived branches originated from lamellipodia. Scale bar indicates 50 μm.*

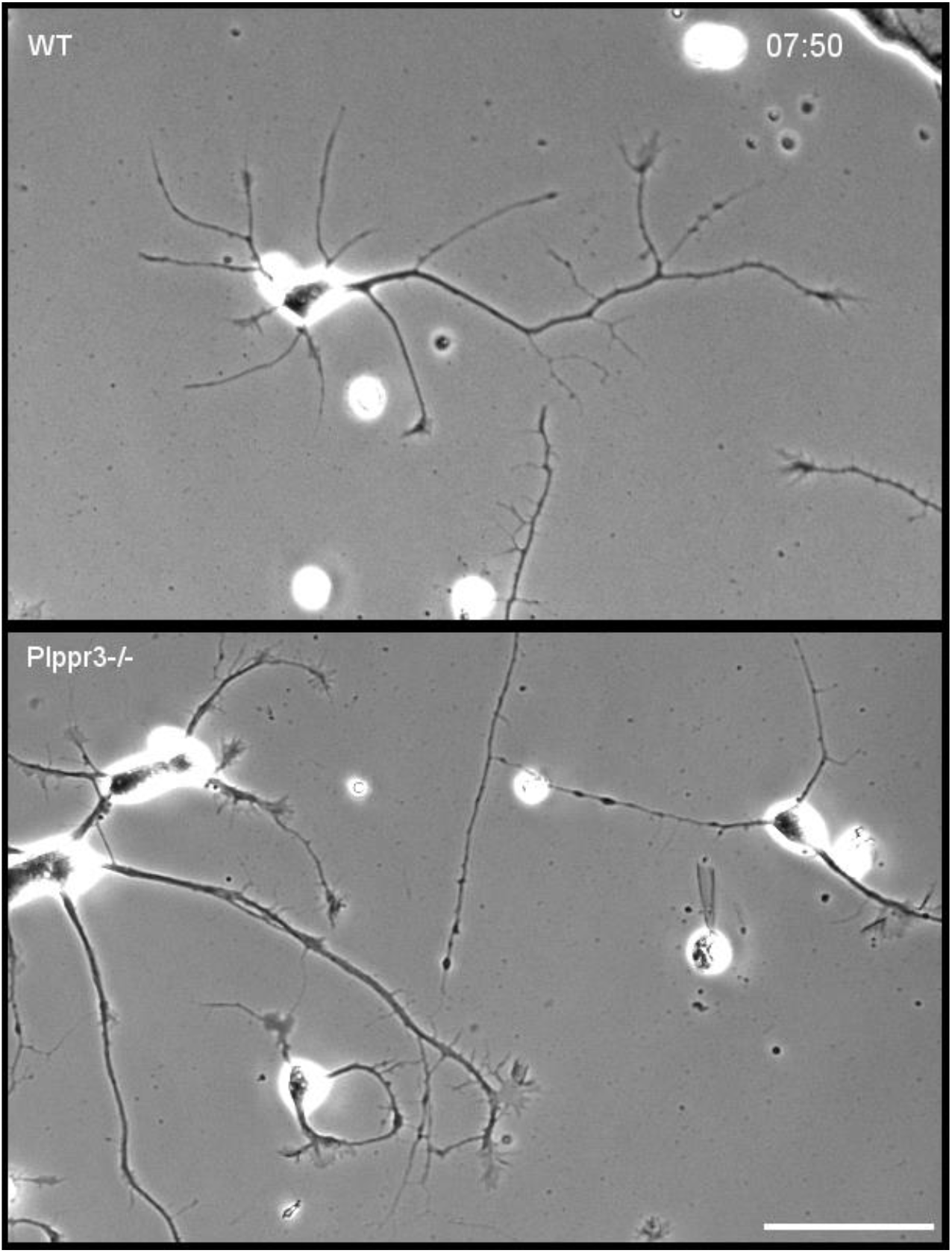
*Individual Timepoint of Supplementary Movie 1 below for illustration*.

**Supplementary figure 1:**
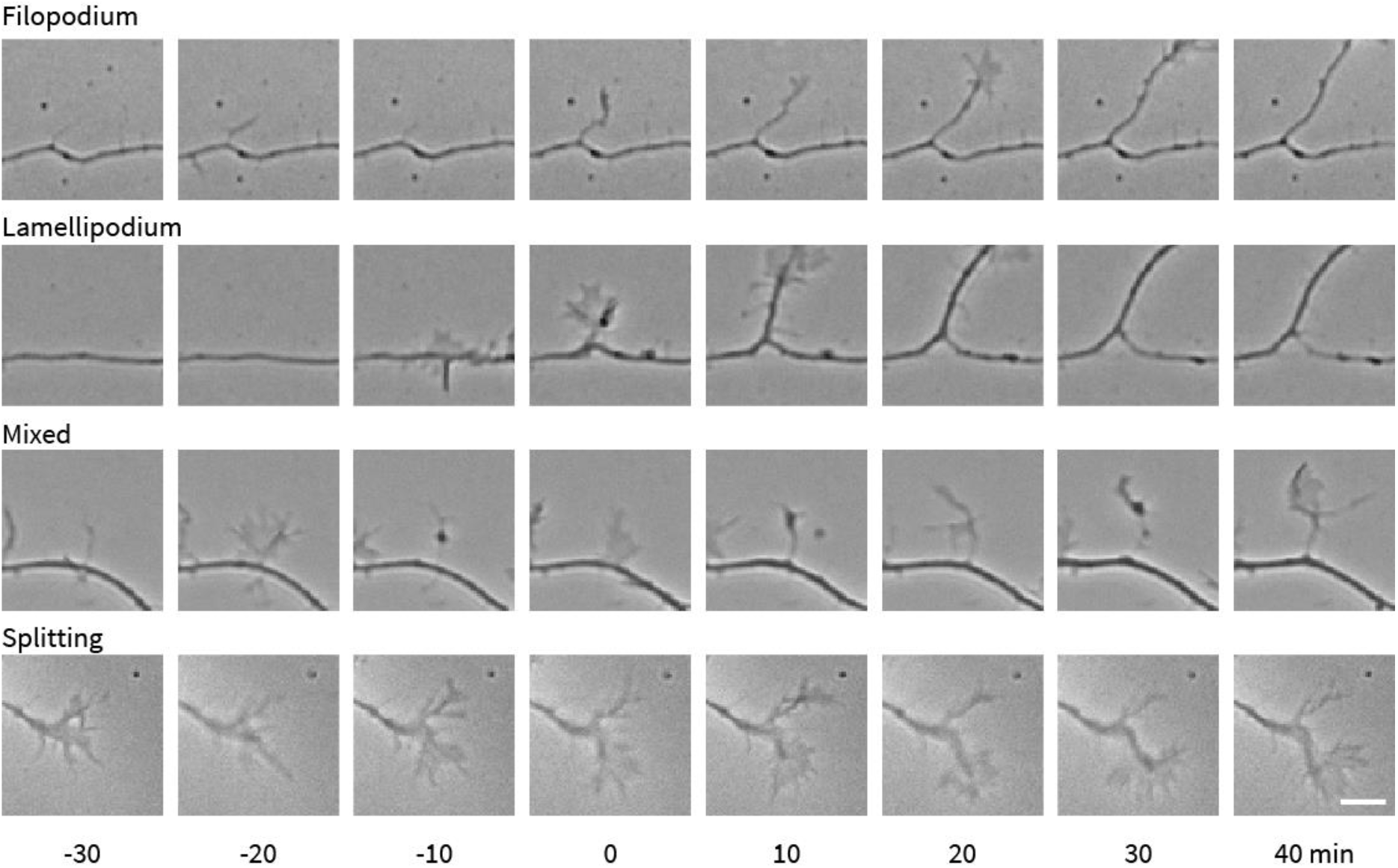
Classification of precursor types by morphology. *and detection of branch emergence and stability over time. Example time series of branch precursor types. 0 indicates the timepoint of branch formation. Scale bar 10 μm.*

**Supplementary figure 2:**
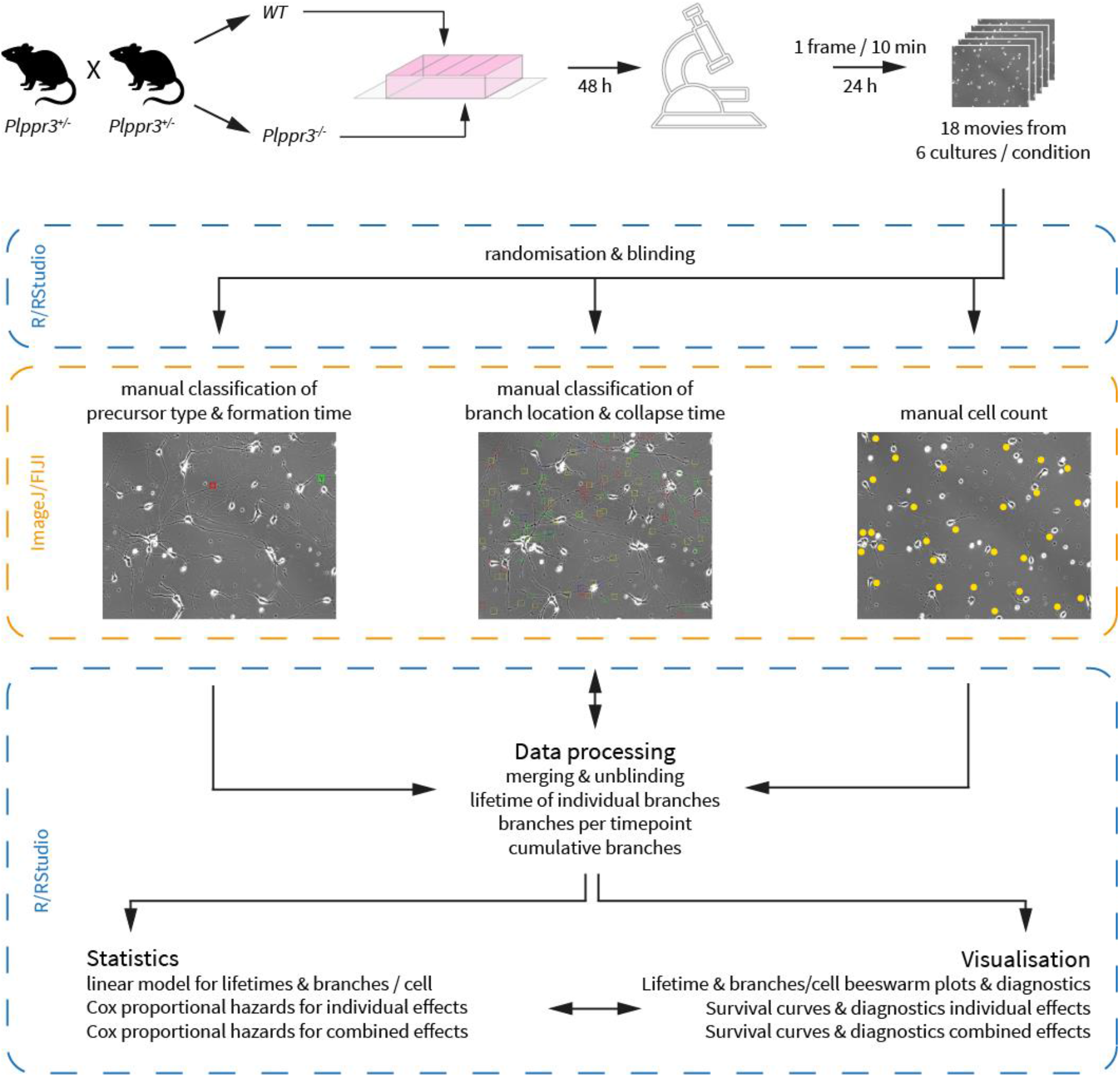
Workflow of experiment and analysis: *Plppr3^−/−^ and WT hippocampal neuron cultures were prepared from littermates of heterozygous Plppr3^+/−^ mice and grown on poly-ornithine & laminin-coated 4-well chamber slides for 48 h. Branching was imaged using phase contrast (Ph2) light microscopy for 24 h with an interval of 10 min. Before manual branch classification in FIJI, each experiment was randomised and blinded to reduce bias in quantification. The analysis consisted of a three-step process, first classifying formation time and precursor type followed by collapse time and location of individual branches. Finally, cells per movie were counted to normalise differences in density between cultures. All subsequent steps of the analysis were performed using R/RStudio and are fully documented at https://github.com/jo-fuchs/Branch-Lifetime-PRG2.*

**Supplementary figure 3:**
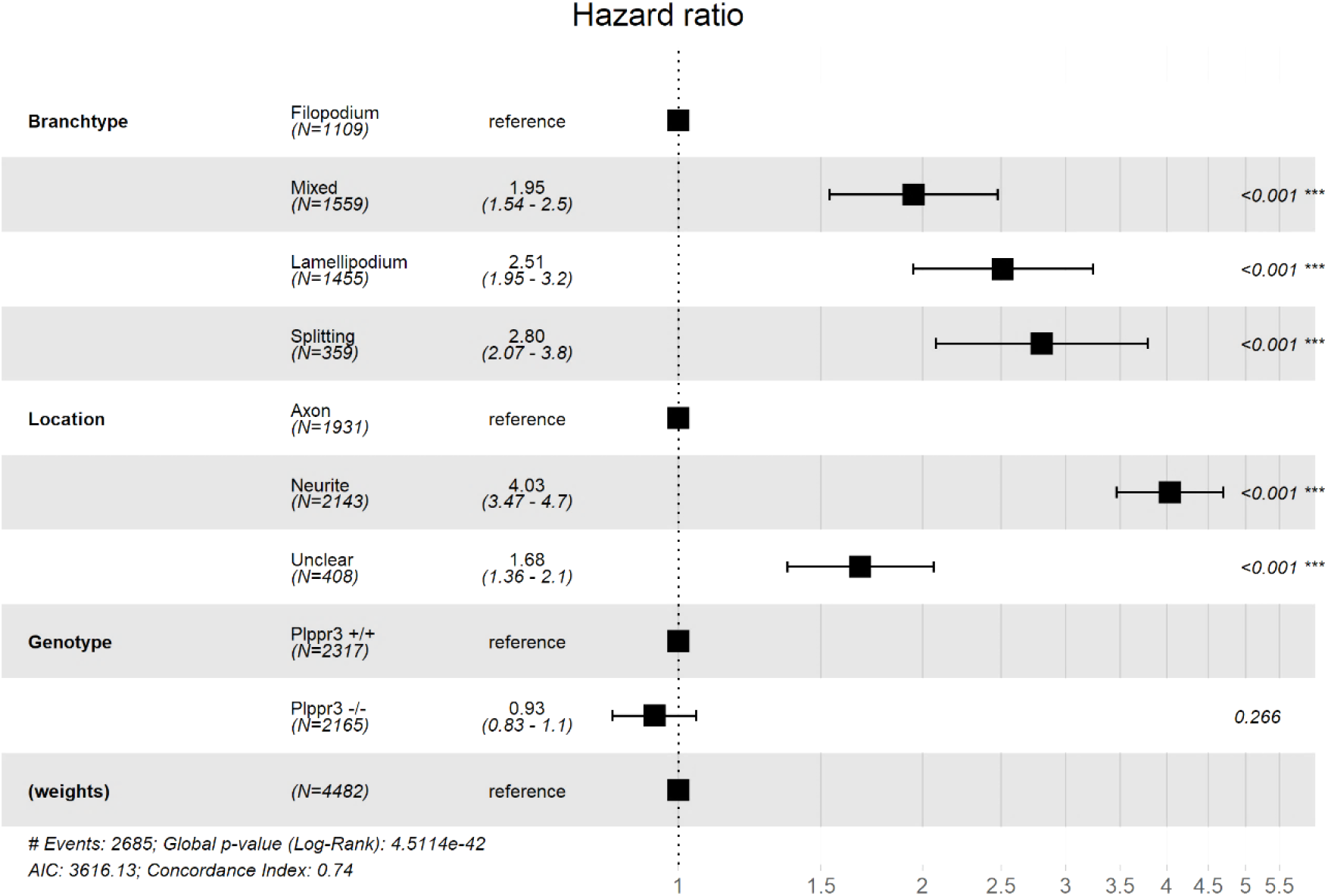
Forest plot of combined Cox Proportional Hazard model used in Figure 2E. *Genotype does not provide any information about branch lifetime after adjusting for precursor type and location.*

## References

Akiyama H, Kamiguchi H. 2010. Phosphatidylinositol 3-Kinase Facilitates Microtubule-dependent Membrane Transport for Neuronal Growth Cone Guidance. J Biol Chem 285:41740–41748. doi:10.1074/jbc.M110.156489

Armijo-Weingart L, Gallo G. 2016. It takes a village to raise a branch: Cellular mechanisms of the initiation of axon collateral branches. Mol Cell Neurosci 84:36–47. doi:10.1016/j.mcn.2017.03.007

Bodakuntla S, Jijumon AS, Villablanca C, Gonzalez-Billault C, Janke C. 2019. Microtubule-Associated Proteins: Structuring the Cytoskeleton. Trends Cell Biol 29:804–819. doi:10.1016/j.tcb.2019.07.004

Brosig A, Fuchs J, Ipek F, Kroon C, Schrötter S, Vadhvani M, Polyzou A, Ledderose J, van Diepen M, Holzhütter HG, Trimbuch T, Gimber N, Schmoranzer J, Lieberam I, Rosenmund C, Spahn C, Scheerer P, Szczepek M, Leondaritis G, Eickholt BJ. 2019. The Axonal Membrane Protein PRG2 Inhibits PTEN and Directs Growth to Branches. Cell Rep 29:2028–2040.e8. doi:10.1016/j.celrep.2019.10.039

Cojoc D, Difato F, Ferrari E, Shahapure RB, Laishram J, Righi M, Di Fabrizio EM, Torre V. 2007. Properties of the Force Exerted by Filopodia and Lamellipodia and the Involvement of Cytoskeletal Components. PLoS One 2:e1072. doi:10.1371/journal.pone.0001072

Conde C, Cáceres A. 2009. Microtubule assembly, organization and dynamics in axons and dendrites. Nat Rev Neurosci 10:319–332. doi:10.1038/nrn2631

Cuntz H, Forstner F, Borst A, Häusser M. 2010. One rule to grow them all: A general theory of neuronal branching and its practical application. PLoS Comput Biol 6. doi:10.1371/journal.pcbi.1000877

Dogterom M, Koenderink GH. 2019. Actin-microtubule crosstalk in cell biology. Nat Rev Mol Cell Biol 20:38–54. doi:10.1038/s41580-018-0067-1

Fink KL, López-Giráldez F, Kim I-J, Strittmatter SM, Cafferty WBJ. 2017. Identification of Intrinsic Axon Growth Modulators for Intact CNS Neurons after Injury. Cell Rep 18:2687–2701. doi:10.1016/j.celrep.2017.02.058

Flynn KC, Pak CW, Shaw AE, Bradke F, Bamburg JR. 2009. Growth cone-like waves transport actin and promote axonogenesis and neurite branching. Dev Neurobiol 69:761–779. doi:10.1002/dneu.20734

Fu X, Brown KJ, Yap CC, Winckler B, Jaiswal JK, Liu JS. 2013. Doublecortin (Dcx) Family Proteins Regulate Filamentous Actin Structure in Developing Neurons. J Neurosci 33:709–721. doi:10.1523/JNEUROSCI.4603-12.2013

Gallo G. 2011. The cytoskeletal and signaling mechanisms of axon collateral branching. Dev Neurobiol 71:201–220. doi:10.1002/dneu.20852

Gibson DA, Ma L. 2011. Developmental regulation of axon branching in the vertebrate nervous system. Development 138:183–195. doi:10.1242/dev.046441

Greenacre M, Primicerio R. 2013. Multivariate Analysis of Ecological Data.

Greenland S, Pearl J, Robins JJM. 1999. Causal Diagrams for Epidemiologic Research. Epidemiology 10:37–48. doi:10.1097/00001648-199901000-00008

Griffin JM, Bradke F. 2020. Therapeutic repair for spinal cord injury: combinatory approaches to address a multifaceted problem. EMBO Mol Med 12:1–29. doi:10.15252/emmm.201911505

Hernán M, Robins J. 2020. Causal Inference: What If. Boca Raton: Chapman & Hall/CRC.

James FC, McCulloch CE. 1990. Multivariate analysis in ecology and systematics: Panacea or Pandora’s box? Annu Rev Ecol Syst 21:129–166. doi:10.1146/annurev.es.21.110190.001021

Janke C, Magiera MM. 2020. The tubulin code and its role in controlling microtubule properties and functions. Nat Rev Mol Cell Biol 21:307–326. doi:10.1038/s41580-020-0214-3

Janson ME, de Dood ME, Dogterom M. 2003. Dynamic instability of microtubules is regulated by force. J Cell Biol 161:1029–1034. doi:10.1083/jcb.200301147

Kakumoto T, Nakata T. 2013. Optogenetic Control of PIP3: PIP3 Is Sufficient to Induce the Actin-Based Active Part of Growth Cones and Is Regulated via Endocytosis. PLoS One 8:1–17. doi:10.1371/journal.pone.0070861

Kalil K, Dent EW. 2014. Branch management: mechanisms of axon branching in the developing vertebrate CNS. Nat Rev Neurosci 15:7–18. doi:10.1038/nrn3650

Kath C, Goni-Oliver P, Müller R, Schultz C, Haucke V, Eickholt B, Schmoranzer J. 2018. PTEN suppresses axon outgrowth by downregulating the level of detyrosinated microtubules. PLoS One 13:1–18. doi:10.1371/journal.pone.0193257

Ketschek A, Gallo G. 2010. Nerve growth factor induces axonal filopodia through localized microdomains of phosphoinositide 3-kinase activity that drive the formation of cytoskeletal precursors to filopodia. J Neurosci 30:12185–12197. doi:10.1523/JNEUROSCI.1740-10.2010

Pearl J. 1995. Causal diagrams for empirical research. Biometrika 82:669–688. doi:10.1093/biomet/82.4.669

Pearl J, Glymour M, Jewell NP. 2016. Causal Inference in Statistics: A Primer. Chichester, England: John Wiley & Sons, Inc.

Pollitt SL, Myers KR, Yoo J, Zheng JQ. 2020. LIM and SH3 protein 1 localizes to the leading edge of protruding lamellipodia and regulates axon development. Mol Biol Cell 31:2718–2732. doi:10.1091/mbc.E20-06-0366

R Core Team. 2020. R: A language and environment for statistical computing. R Found Stat Comput Vienna, Austria. http://www.r-project.org/index.html

Rohrer JM. 2018. Thinking Clearly About Correlations and Causation: Graphical Causal Models for Observational Data. Adv Methods Pract Psychol Sci 1:27–42. doi:10.1177/2515245917745629

Schindelin J, Arganda-Carreras I, Frise E, Kaynig V, Longair M, Pietzsch T, Preibisch S, Rueden C, Saalfeld S, Schmid B, Tinevez J-Y, White DJ, Hartenstein V, Eliceiri K, Tomancak P, Cardona A. 2012. Fiji: an open-source platform for biological-image analysis. Nat Methods 9:676–682. doi:10.1038/nmeth.2019

Spillane M, Ketschek A, Merianda TT, Twiss JL, Gallo G. 2013. Mitochondria Coordinate Sites of Axon Branching through Localized Intra-axonal Protein Synthesis. Cell Rep 5:1564–1575. doi:10.1016/j.celrep.2013.11.022

Suttorp MM, Siegerink B, Jager KJ, Zoccali C, Dekker FW. 2015. Graphical presentation of confounding in directed acyclic graphs. Nephrol Dial Transplant 30:1418–1423. doi:10.1093/ndt/gfu325

Tedeschi A, Dupraz S, Curcio M, Laskowski CJ, Schaffran B, Flynn KC, Santos TE, Stern S, Hilton BJ, Larson MJE, Gurniak CB, Witke W, Bradke F. 2019. ADF/Cofilin-Mediated Actin Turnover Promotes Axon Regeneration in the Adult CNS. Neuron 103:1073–1085.e6. doi:10.1016/j.neuron.2019.07.007

Tymanskyj SR, Yang B, Falnikar A, Lepore AC, Ma L. 2017. MAP7 regulates axon collateral branch development in dorsal root ganglion neurons. J Neurosci 37:1648–1661. doi:10.1523/JNEUROSCI.3260-16.2017

Willige D, Hummel JJ, Alkemade C, Kahn OI, Au FK, Qi RZ, Dogterom M, Koenderink GH, Hoogenraad CC, Akhmanova A. 2019. Cytolinker Gas2L1 regulates axon morphology through microtubule‐modulated actin stabilization. EMBO Rep 20:1–20. doi:10.15252/embr.201947732

Winans AM, Collins SR, Meyer T. 2016. Waves of actin and microtubule polymerization drive microtubule-based transport and neurite growth before single axon formation. Elife 5:1–22. doi:10.7554/elife.12387

Withers GS, Wallace CS. 2020. Transient lamellipodia predict sites of dendritic branch formation in hippocampal neurons. Cell Tissue Res. doi:10.1007/s00441-020-03194-w

Yu W, Qiang L, Solowska JM, Karabay A, Korulu S, Baas PW. 2008. The Microtubule-severing Proteins Spastin and Katanin Participate Differently in the Formation of Axonal Branches. Mol Biol Cell 19:1485–1498. doi:10.1091/mbc.e07-09-0878

